# GM-CSF differentiation of human monocytes stabilizes macrophage state via oxidative signaling

**DOI:** 10.1101/2020.09.29.318352

**Authors:** Christopher Y Itoh, Cal Gunnarsson, Gregory H Babunovic, Armel Nibasumba, Ngomu Akeem Akilimali, Marc H Wadsworth, Travis K Hughes, Sydney L Solomon, Brian Hie, Bonnie Berger, Alex K Shalek, Sarah M Fortune, Bryan D Bryson

## Abstract

Macrophages are central mediators of immunity that integrate diverse signals derived from differentiation cues, tissue location, and disease. Controlling macrophage state and function is an appealing therapeutic objective across many diseases including cancer, atherosclerosis, and tuberculosis. Despite the growing appreciation for the *in vivo* complexity of macrophage state, existing *in vitro* models of human monocyte-derived macrophages have used a limited number of individual perturbations to explore the complex phenotypic space that macrophages can occupy. Here, we leverage a tiered differentiation, activation, and stimulation strategy to generate libraries of *in vitro* monocyte-derived macrophages and examine the *in vitro* state space of macrophage function using high-dimensional technologies. Our tiered experimental approach further revealed a striking relationship between GM-CSF differentiation and IL-10 production. Cells that were differentiated with GM-CSF produced very low or undetectable levels of IL-10 independent of activation or stimulation condition. To nominate candidate regulators of this IL-10 response, we leverage unbiased single-cell mRNA sequencing to identify transcriptional networks associated with GM-CSF-derived cells. Using these data, we identify oxidative signaling pathways as upregulated in GM-CSF derived cells and demonstrate that scavenging of oxidative radicals can enhance IL-10 production in these cells. Collectively, these data underscore the complexity of monocyte-derived macrophage state over time and highlight a dominant role for GM-CSF in tuning macrophage inflammatory phenotype, metabolic state, and plasticity.

## Introduction

Understanding how macrophages interpret diverse signal inputs to dictate their state and function is an actively evolving field. While previous nomenclature to describe macrophage state adopted an M1 or M2 naming convention to parallel elements of Th1 and Th2 immunity, new naming conventions integrate additional dimensions of macrophage diversity including differentiation or activation cytokine and signal duration (Murray et al., 2014). This naming convention recognizes that distinct cues can drive specific cytokine-associated programs in addition to generic inflammatory or anti-inflammatory programs that are conserved across diverse inputs. As we continue to identify additional cues that redirect macrophage function, it is important to understand how these signals reinforce existing macrophage states or perturb them. Due to this evolving nomenclature, determining macrophage state using standard tools such as flow cytometry with canonical macrophage markers remains challenging as these markers are often expressed independent of macrophage state.

Recent murine studies highlight the complex and exquisite trajectories monocytes traverse as they differentiate into tissue-resident macrophages. In fact, in many tissues, monocyte-derived macrophages serve as the dominant contributor to the steady-state macrophage pool (Liu et al., 2019). These *in vivo* studies have been utilized to highlight features of macrophage plasticity and identify the signals that drive tissue-resident macrophage identity. For example, Lavin and colleagues demonstrated that tissue-resident macrophages can be transferred to new tissues and adopt features of the new microenvironment (Lavin et al., 2014). They demonstrated that peritoneal macrophages could be transferred intratracheally to the alveolar cavity of recipient mice and adopt features of the alveolar macrophages as evidenced by transcriptional profiling. These studies and countless others have been utilized to demonstrate features of macrophage phenotypic plasticity.

In contrast to the study of experimentally available murine tissue-resident macrophages, studies of human macrophages rely mostly on monocyte-derived macrophages differentiated from peripheral blood mononuclear cells of human donors. To differentiate these monocytes into macrophages, common protocols supplement culture media with either macrophage colony-stimulating factor (M-CSF) or granulocyte macrophage colony-stimulating factor (GM-CSF). Sander and colleagues utilized transcriptional profiling to demonstrate that human monocytes differentiated with M-CSF or GM-CSF resembled macrophages isolated directly from human ascites thus underscoring their experimental utility (Sander et al., 2017). Additionally, the authors demonstrated that monocytes integrate simultaneous signals from IL-4 and GM-CSF distinguishing them from monocytes differentiated with GM-CSF alone and suggest that varying the exposure time of IL-4 tunes the functional identity of the cells. Taken together, these previous studies highlight the complexity of macrophage behavior and their modes of signal integration.

To build upon these previous studies and examine how signal identity and duration influence human macrophage functional state, we established an *in vitro* experimental platform to create libraries of monocyte-derived macrophages. Using a combination of flow cytometry, cytokine release assays, and single-cell RNA sequencing (scRNAseq), we demonstrate that GM-CSF differentiation of monocytes abolishes the capacity of these cells to produce IL-10 in response to TLR ligands. Moreover, we were unable to identify a cytokine or small molecule that was capable of restoring IL-10 production in these cells following differentiation. This finding provides an important update to the long-held model of macrophage plasticity. Using scRNAseq, we overcome existing challenges in macrophage characterization and identify novel markers of macrophage state that permit discrimination of macrophages on the basis of differentiation cytokine. We further leverage these scRNAseq results and nominate oxidative signaling as a driver of reduced IL-10 production in macrophages differentiated by GM-CSF which we then confirmed in our *in vitro* system. Taken together, these data emphasize the complexity of macrophage functional properties and suggest a dominant role for GM-CSF in modulating macrophage state.

## Results

### Generation of libraries of *in vitro* monocyte-derived macrophages provides flexible platform for interrogating macrophage state and function

Human monocytes differentiated with M-CSF or GM-CSF are widely used as *in vitro* models for studying macrophage behavior (Lacey et al., 2012; Sander et al., 2017; Xue et al., 2014). To examine the contribution of these diverse axes, we generated libraries of *in vitro* monocyte-derived macrophages from CD14+ monocytes isolated from healthy human donors using a systematic framework for perturbation and characterization. CD14+ monocytes were first differentiated for 3 days with either GM-CSF or M-CSF to generate MO-GMCSF or MO-MCSF. To generate additional diversity in our *in vitro* cultures, on day 3 of differentiation, we supplemented our differentiation media with an additional activation stimulus (vehicle, LPS, IFN-*γ*, IL-4, IL-6, and IL-10) ultimately generating a total of 12 macrophage populations (**Figure 1A**) (Roussel et al., 2017).

**Figure 1:**
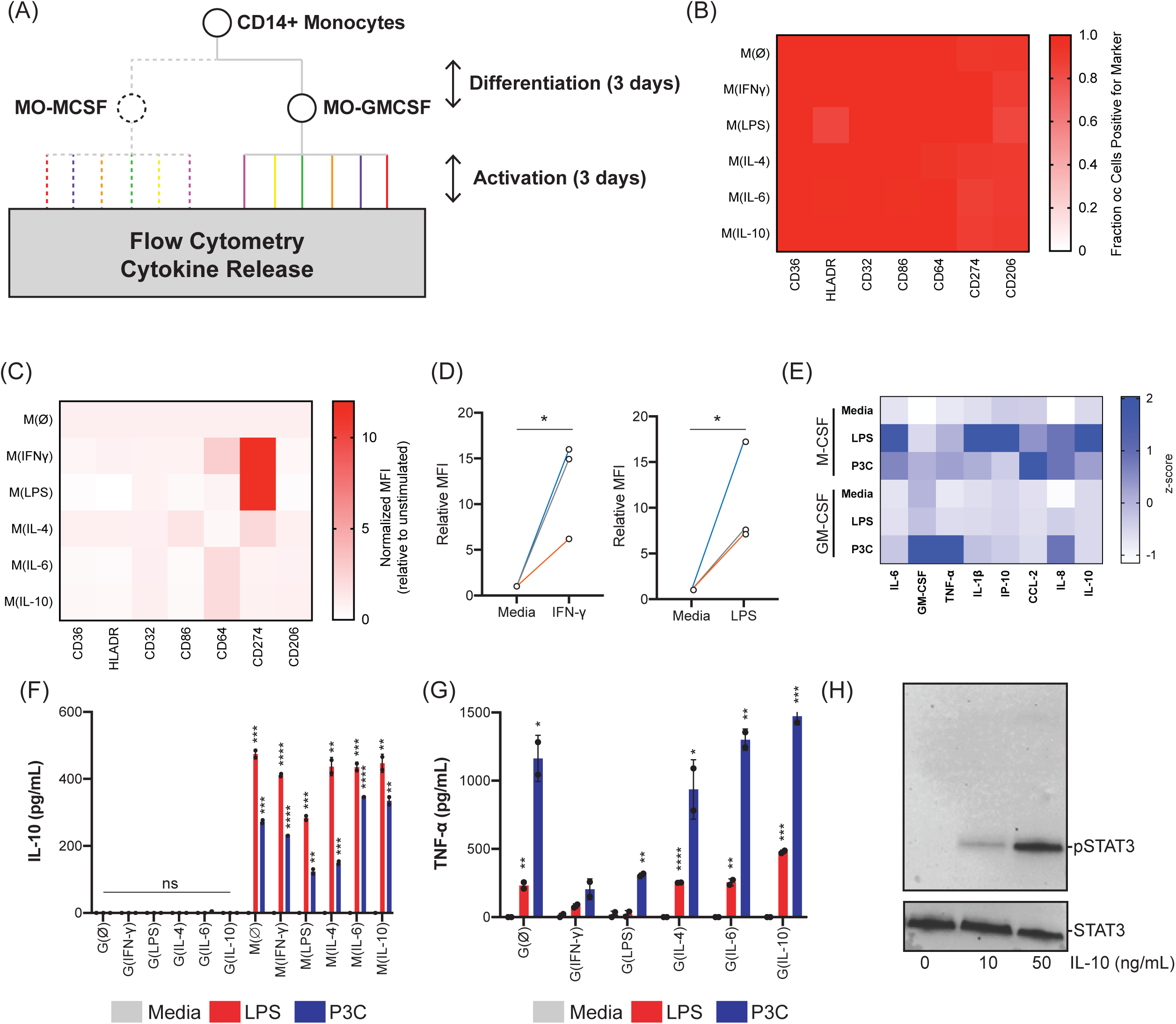
Multi-dimensional profiling of human monocyte-derived macrophages identifies differentiation and activation-mediated phenotypes. (A) Methodological workflow for generation of macrophage libraries from CD14+ monocytes from healthy donors. Monocytes were differentiated with M-CSF or GM-CSF for 3 days (differentiation only phase) prior to the addition of additional ligands (PBS, LPS, IFN-*γ*, IL-4, IL-6, and IL-10) for an additional 3 days (activation phase). After 3 days in the activation phase, macrophages were washed with PBS, detached, counted, and re-plated equally to characterize macrophage phenotype. (B) Analysis of macrophage surface marker expression using a digital gating strategy to identify the percentage of macrophages expressing a given marker (MO-MCSF cells). Representative of 3 donors. (C) Analysis of macrophage surface marker expression using an analog quantification strategy (mean fluorescence intensity, MFI) to quantify the abundance of a given marker on the macrophage surface (MO-MCSF cells). Representative of 3 donors. (D) Quantification of CD274 across multiple donors reveals upregulation of CD274 following IFN-*γ* or LPS stimulation (MO-MCSF cells). Each colored line is an individual donor. Comparisons made using a ratio paired t test. (*, p < 0.05). (E) Z-score of cytokine secretion screen in MO-MCSF or MO-GMCSF cells stimulated with media, 100 ng/mL LPS or 100 ng/mL Pam3Csk4 (P3C). (F) Secretion of IL-10 across all 12 macrophage populations following 100 ng/mL LPS or 100 ng/mL P3C stimulation. (****, p < 0.0001, ***, p < 0.001, **, p < 0.01, ns = not significant). Representative of 3 donors. (G) Secretion of TNF*α* across all 6 MO-GMCSF populations following 100 ng/L LPS or 100 ng/mL P3C stimulation. (****, p < 0.0001, ***, p < 0.001, **, p < 0.01, *, p <0.05, ns = not significant). Representative of 3 donors. (H) Phosphorylation of STAT3 in MO-GMCSF cells following IL-10 stimulation.

We first utilized flow cytometry to characterize macrophage state by selecting commonly used macrophage surface markers from previous studies (Roussel et al., 2017). We compared surface expression of CD32, CD36, CD64, CD86, CD206, CD274, and HLA-DR focusing first on the differences in MO-MCSF and MO-GMCSF libraries separately. We used two metrics to characterize surface expression of these markers across our samples: (1) percentage of cells expressing any given marker relative to an isotype control and (2) the mean fluorescence intensity (MFI) of the surface markers. We sought to examine if any of the markers in our panel demonstrated digital expression among the macrophages generated in our *in vitro* libraries. In contrast to lymphocytes where several markers exist to distinguish subpopulations using a binary gating strategy, none of the macrophage markers used in this panel were capable of discriminating between the different macrophage populations (**Figure 1B, Supplemental Figure 1)** (DuPage and Bluestone, 2016; Murray et al., 2014). As previously observed by others, our results displayed differences in the level of expression of various markers across all macrophage types (**Figure 1C, Supplemental Figure 2**) (Schulz et al., 2019). Specifically, we observed CD274 consistently upregulated in cells stimulated with IFN-*γ* or LPS in both MO-MCSF and MO-GMCSF cells. **(Figure 1D (MO-MCSF), Supplemental Figure 3 (MO-GMCSF))**.

As a complementary approach to flow cytometry, we next characterized our macrophage library by examining cytokine release in response to purified TLR ligands. We initially conducted a multiplex cytokine screen focusing on the differences between MO-MCSF and MO-GMCSF in response to lipopolysaccharide (LPS), a TLR4 agonist and Pam3CSK4 (PAM), a TLR1/2 agonist, for 24 hours prior to cytokine analysis. We measured the release of several cytokines, including IL-1*β*, IL-6, and IL-10. Comparing MO-MCSF and MO-GMCSF cells, we observed a marked reduction in the production of IL-10 and CCL-2 in MO-GMCSF cells in response to both TLR ligands, consistent with previous reports comparing these cells (**Figure 1E**) (Lacey et al., 2012; Sander et al., 2017). We next extended these measurements to all twelve populations of MO-MCSF and MO-GMCSF cells. We focused on the production of TNF*α*, IL-6, and IL-10 given the dynamic range of their response to TLR stimulation. By enzyme-linked immunosorbent assay (ELISA), we were able to validate our observation of decreased production of IL-10 in GM-CSF derived cells. Notably, we observed levels of IL-10 production that were near or below the ELISA limit of detection across all MO-GMCSF cells independent of TLR stimulation or activation cue (**Figure 1F**). This was in contrast to M-CSF derived macrophages which retained their ability to produce IL-10. MO-GMCSF cells, despite lacking an IL-10 response, retained their ability to respond to TLR ligands and produce cytokines as indicated by production of TNF*α* (**Figure 1G**). We sought to recapitulate these findings using intracellular cytokine staining; however, we were unable to find a antibody clone that demonstrated robust differences between unstimulated and stimulated MO-MCSF cells (**Supplemental Figure 4**).

Since our cytokine secretion results pointed to an altered IL-10 signaling network in MO-GMCSF cells, we next sought to determine if MO-GMCSF cells were muted in both their IL-10 production and responsiveness or production alone. To assess the capacity of MO-GMCSF cells to respond to IL-10, we monitored the phosphorylation status of STAT3, a well-characterized protein downstream of IL10R, following stimulation with IL-10 (Niemand et al., 2003). We observed a dose-dependent increase in phosphorylated STAT3 relative to total STAT3 in MO-GMCSF cells indicating that GM-CSF differentiation does not abrogate IL10R signaling (**Figure 1H)**. Collectively, these data demonstrate how macrophage differentiation cues can precisely fine-tune macrophage state and illustrate the utility of diverse modalities to characterize macrophage phenotypes.

### scRNAseq reveals novel markers of the MO-GMCSF state

Given the phenotypic differences observed between MO-MCSF and MO-GMCSF cells, we employed scRNAseq to identify additional markers capable of distinguishing between these two macrophage states. To robustly interrogate the transcriptomes of thousands of cells we used Seq-Well, a nanowell platform for high-throughput scRNAseq (Gierahn et al., 2017). This produced 1,742 high-quality MO-GMCSF and 4,803 high-quality MO-MCSF cellular transcriptomes from 2 donors (**Supplementary Figure 5**). We first examined the expression of canonical markers of macrophage state to characterize their ability to discriminate between MO-MCSF and MO-GMCSF cells. A subset of canonical marker genes achieved statistically significant differential expression (CD68, CD86, MRC1 (CD206); however, the magnitude of their transcriptional difference was minimal (**Figure 2A, Supplementary Table 1**). Other canonical markers either displayed no significant differences or had low capture rates. We next examined the expression of all genes measured by scRNAseq. Using a Wilcoxon rank sums test, we identified 178 significantly upregulated and 137 significantly downregulated genes in MO-GMCSF cells relative to MO-MCSF cells (**Figure 2B**). We identified several novel marker genes of MO-MCSF and MO-GMCSF cells. We identified CHI3L1 and MGLL as markers of MO-MCSF and TXN and ALOX5AP as markers of MO-GMCSF (**Figure 2C**). These genes included CHI3L1 and MGLL as markers of MO-MCSF and TXN and ALOX5AP as markers of MO-GMCSF. To gain an unbiased view of the transcriptional programs differentially active in MO-MCSF and MO-GMCSF cells, we next identified Gene Ontology Molecular Function (GO MF) terms enriched in the set of marker genes for each state. This analysis revealed MO-GMCSF cells were marked by transcriptional programs related to redox signaling, as well as antigen presentation (**Figure 2D, Supplementary Table 2**).

**Figure 2:**
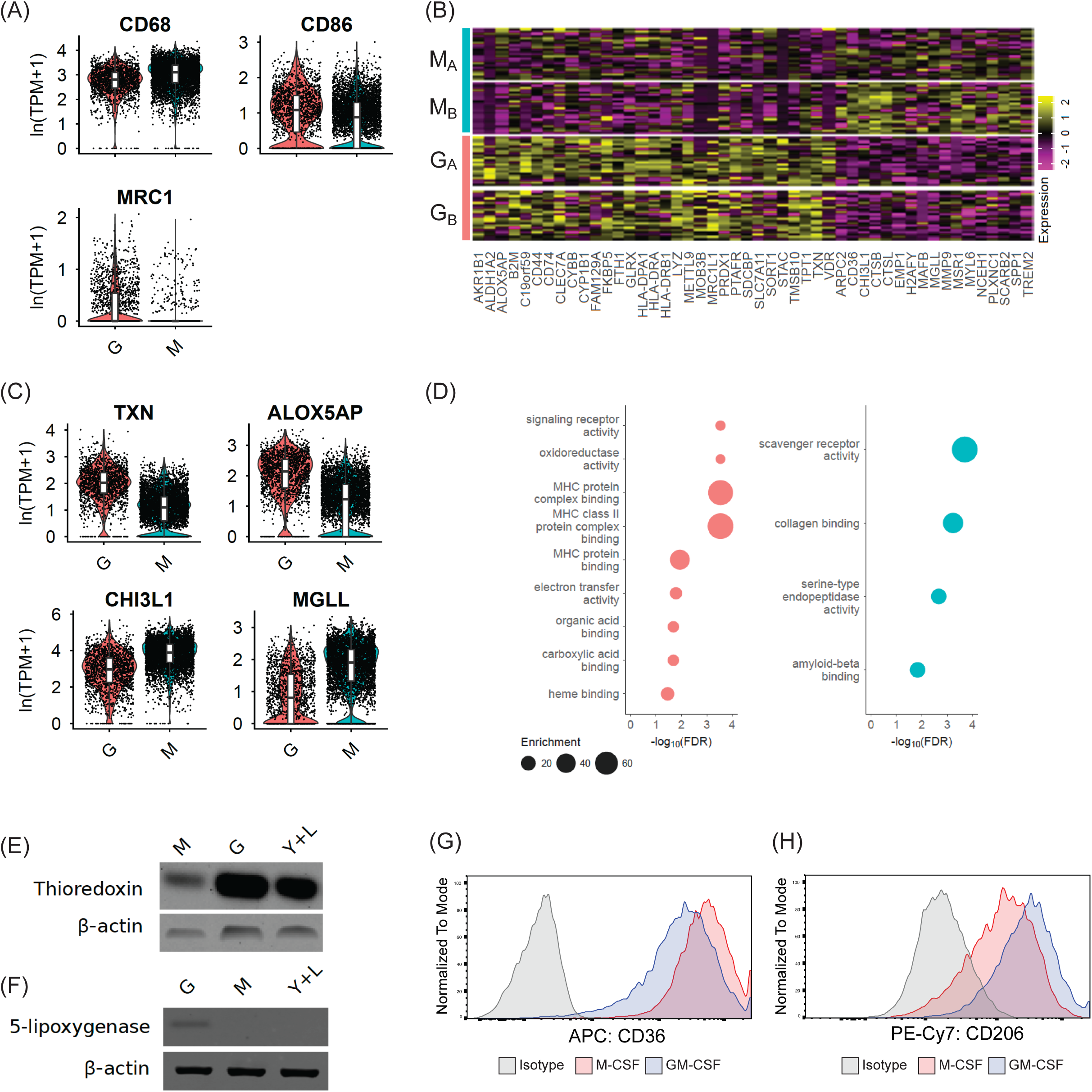
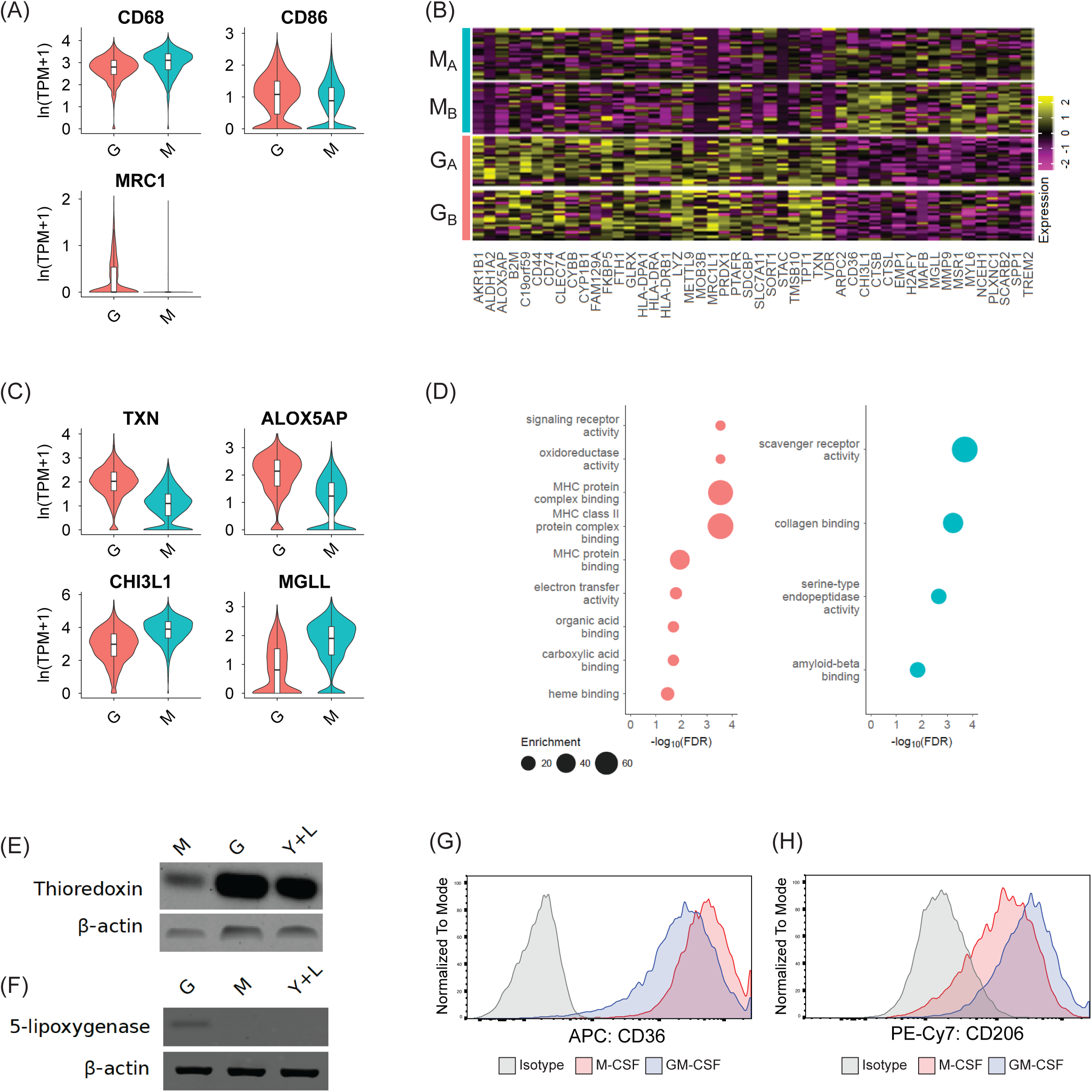
Single cell RNA sequencing reveals characteristics of macrophage state. (A) Log-normalized expression of canonical marker genes in 1,742 MO-GMCSF and 4,803 MO-MCSF from 2 donors (N = 399 MO-GMCSF- and 1,941 MO-M-CSF donor 1; 1,343 MO-GMCSF- and 2,862 MO-MCSF-treated cells, donor 2). Center line, median; box limits, first and third quartile; whiskers, +/- 1.5 interquartile range; dots, individual cells. G = MO-GMCSF-treated cells; M = MO-MCSF-treated cells. (B) Standard scaled expression of marker genes identified in 250 randomly subsampled cells from each donor and condition. M_A_ = M-CSF-treated, donor A; M_B_ = M-CSF-treated, donor 2; G_A_ = GM-CSF-treated, donor A; G_B_ = GM-CSF-treated, donor B. (C) Log-normalized expression of top marker genes identified in (B) across all cells. (AUC, TXN: 0.845, ALOX5AP: 0.808, CHI3L1: 0.222, MGLL: 0.225) (D) Significantly enriched gene ontology terms in marker gene set for MO-GMCSF-cells (left) and M-CSF-treated cells (right), determined by hypergeometric test after filtering by semantic similarity (Benjamini-Hochberg-corrected p-value < 0.05; semantic similarity threshold = 0.5). GO term enrichment in the marker set was calculated relative to enrichment in the background set of genes captured in all sequencing runs with >100 total counts across runs. Western blotting of cell lysates treated with 50 ng/mL M-CSF, 50 ng/mL GM-CSF, or 10 ng/mL each of IFN-*γ* and LPS (Y+L) for 24 h for thioredoxin (E) and 5-lipoxygenase (F). Beta-actin was used as a loading control. Representative of 4 donors. (G) Analysis of CD36 expression by flow cytometry in MO-MCSF or MO-GMCSF cells. Representative of 3 donors. (H) Analysis of CD206 expression by flow cytometry in MO-MCSF or MO-GMCSF cells. Representative of 3 donors.

Using scRNAseq, we found several novel markers that distinguished MO-MCSF and MO-GMCSF cells. The gene encoding thioredoxin (*TXN*) was a robust marker of MO-GMCSF cells by an AUC classifier (AUC = 0.85). Thioredoxin is an antioxidant that is involved in many different signaling pathways via redox control (Lee et al., 2013). Previous studies in human macrophage-like cell lines have demonstrated that thioredoxin is highly expressed in response to the combination of LPS and IFN-*γ*, a well-characterized inducer of an inflammatory phenotype (Plugis et al., 2018). Other studies have suggested that MO-GMCSF cells may serve as another model of inflammatory macrophages (Fleetwood et al., 2007; Hamilton, 2019). We hypothesized that the induction of thioredoxin may be conserved in both MO-MCSF treated with LPS+IFN-*γ* and MO-GMCSF cells. Consistent with the scRNAseq data and our hypothesis, thioredoxin protein was highly expressed in MO-GMCSF, as well as M(LPS+IFN-*γ*), compared to MO-MCSF cells (**Fig 2E**). We next sought to validate an additional marker of MO-GMCSF, 5-lipoxygenase accessory protein (ALOX5AP), which is essential for the production of inflammatory lipids such as prostaglandin E2. Again, consistent with our scRNAseq data, ALOX5AP was highly expressed in MO-GMCSF cells relative to MO-MCSF cells; however, ALOX5AP was not highly expressed in M(LPS+IFN-*γ*), suggesting distinct regulatory axes contributing to expression of these marker genes (**Figure 2F**). These results are consistent with previous work demonstrating altered expression of ALOX5AP and dichotomous production of inflammatory and pro-resolving lipids (Werz et al., 2018). Further supporting our scRNAseq results, we also observed enhanced expression of CD36 on MO-MCSF cells and increased expression of CD206 on MO-GMCSF cells by flow cytometry (**Figure 2G, Figure 2H**). Taken together, these results identify novel markers that can more definitively characterize macrophage state and distinguish common transcriptional programs that are active across macrophage culture systems.

### GM-CSF mediates a durable switch in IL-10 production

We reasoned that the immunophenotyping analyses had provided us with markers that we could leverage to interrogate phenotypic transitions between macrophage state. Macrophages have long been described as cells that demonstrate exquisite phenotypic plasticity; however, our IL-10 secretion results suggested that following GM-CSF differentiation, several cytokines were unable to enhance TLR-mediated IL-10 production to levels above the detection limit (Biswas and Mantovani, 2010; Liu et al., 2020; Locati et al., 2020).

Using IL-10 production in response to TLR stimulation as a measure of macrophage state, we sought to interrogate the stability of the MO-MCSF and MO-GMCSF phenotype. We first sought to examine the expression of M-CSF and GM-CSF receptors on the surface of MO-MCSF and MO-GMCSF cells reasoning that we might be able to utilize the opposing cytokine to shift macrophage state. We performed flow cytometry on MO-MCSF or MO-GMCSF cells following 6 days of differentiation. We detected surface expression of GM-CSF (CD116) and M-CSF (CD115) receptors on the surface of MO-MCSF and MO-GMCSF cells (**Figure 3A**). We next sought to examine how treatment of MO-MCSF and MO-GMCSF cells with the opposing cytokine would modulate IL-10 production. Monocytes were differentiated in the presence of M-CSF or GM-CSF for 6 days, washed, and then cultured in media containing 50 ng/mL M-CSF, media containing 50 ng/mL GM-CSF, or media alone for an additional 3 days. After this second phase of culture, cells were washed and then stimulated with LPS for 24 hours prior to analysis of IL-10 secretion (**Figure 3B**). As expected from our previous observations, MO-MCSF cells that received no cytokine or M-CSF during the second media phase produced IL-10 in response to LPS. In contrast, MO-MCSF cells treated with GM-CSF during the second media phase produced less IL-10 in response to LPS (**Figure 3C**). Strikingly, MO-GMCSF cells independent of their second media phase did not produce IL-10 at levels above the limit of detection in response to LPS stimulation similar to our results observed with IFN-*γ*, IL-4, IL-6, IL-10, and LPS.

**Figure 3:**
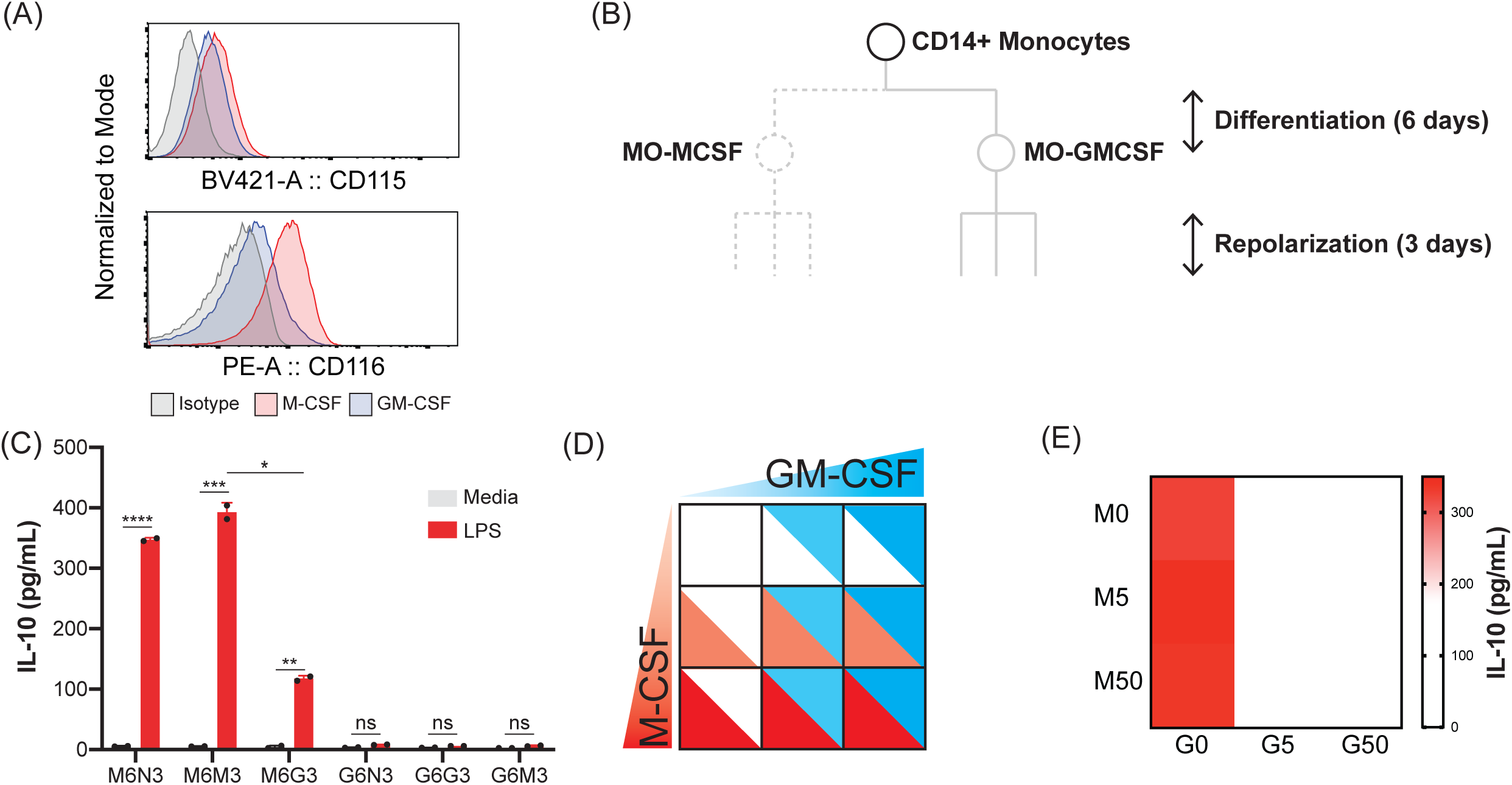
Combinatorial treatment of monocytes or MO-MCSF or MO-GMCSF cells with GM-CSF or M-CSF reveals unexpected features of the GM-CSF-IL-10 circuit. (A) Analysis of GM-CSF (CD116) or M-CSF (CD115) receptor surface expression by flow cytometry in MO-MCSF or MO-GMCSF cells. Representative of 3 donors. (B) Design of IL-10 phenotypic stability experiment in MO-MCSF or MO-GMCSF cells. (C) Secretion of IL-10 across polarized or repolarized MO-MCSF and MO-GMCSF cells following 100 ng/mL LPS stimulation. (****, p < 0.0001, ***, p < 0.001, **, p < 0.01, ns = not significant). Representative of 3 donors. (D) Design of GM-CSF/M-CSF titration checkerboard experiment. (E) Secretion of IL-10 across monocytes differentiated with various combinations of GM-CSF or M-CSF following 100 ng/mL LPS stimulation. Representative of 3 donors.

While macrophages may encounter asynchronous stimuli as modeled by temporally staggering the addition of cytokines, we reasoned that monocytes upon recruitment to tissues may simultaneously encounter combinations of cytokines such as M-CSF and GM-CSF that may act synergistically to drive the functional state of the resulting macrophages. We next sought to examine whether the monocyte response to GM-CSF or M-CSF differentiation could be modeled as an additive response. To examine these dynamics, we generated a matrix of combinations of M-CSF and GM-CSF ranging from 0 ng/mL to 50 ng/mL in ten-fold increments (**Figure 3D, Figure 3E**). Cytokines were premixed and arrayed prior to the addition of monocytes to eliminate any temporal asynchrony in cytokine exposure. Monocytes were differentiated for 6 days, washed, and then stimulated with LPS. IL-10 production was monitored by ELISA. Monocytes cultured for 6 days in the absence of exogenous cytokines on tissue-culture treated plasticware did not survive. In the absence of GM-CSF, we observed a dose-dependent increase in LPS-mediated IL-10 release as we increased the concentration of M-CSF in the differentiation media. Holding the M-CSF concentration at 0 ng/mL, we observed no increase in LPS-mediated IL-10 production as the concentration of GM-CSF increased. Intriguingly, nearly all combinations of M-CSF and GM-CSF resulted in low or undetectable levels of IL-10 in response to LPS suggesting a non-additive hypersensitivity to GM-CSF in driving macrophage state as previously observed at the transcriptional level (Lacey et al., 2012). Taken together, these data highlight the potency of GM-CSF in modulating macrophage state and underscore the importance of understanding the additive and non-additive gene and protein expression relationships in myeloid cells.

### Disruption of elements of the canonical GM-CSF signaling network does not enhance IL-10 production in MO-GMCSF cells but scavenging of ROS does

Our tiered macrophage differentiation strategy did not identify a cytokine capable of restoring IL-10 production in MO-GMCSF cells. As an alternative strategy to mechanistically dissect the drivers of muted IL-10 production in MO-GMCSF cells, we turned to previously established models for modifying either macrophage phenotypic stability or IL-10 production.

Previous studies in murine and human macrophages have demonstrated that macrophages adopt semi-stable phenotypes as demonstrated by surface marker and marker gene expression (Van den Bossche et al., 2016). In these studies, the authors implicate mitochondrial dysfunction mediated by inducible nitric oxide synthase (iNOS) as a driver of limited M1 to M2 repolarization in murine macrophages. We sought to test the hypothesis that iNOS may play a role in the limited plasticity observed in MO-GMCSF cells recognizing that expression of iNOS in human macrophages has been widely debated (Bogdan, 2001; Schneemann and Schoedon, 2002). Nevertheless, we sought to reproduce the experimental model employed previously to examine functional stability in MO-GMCSF macrophages. Human monocytes were pre-treated with 1400W, an iNOS inhibitor, or DMSO for one hour prior to the addition of M-CSF or GM-CSF. After 6 days, the cells were washed and stimulated with LPS. Production of IL-10 was observed in all MO-MCSF cells; however, IL-10 remained undetectable in MO-GMCSF cells across all conditions (**Figure 4A**).

**Figure 4:**
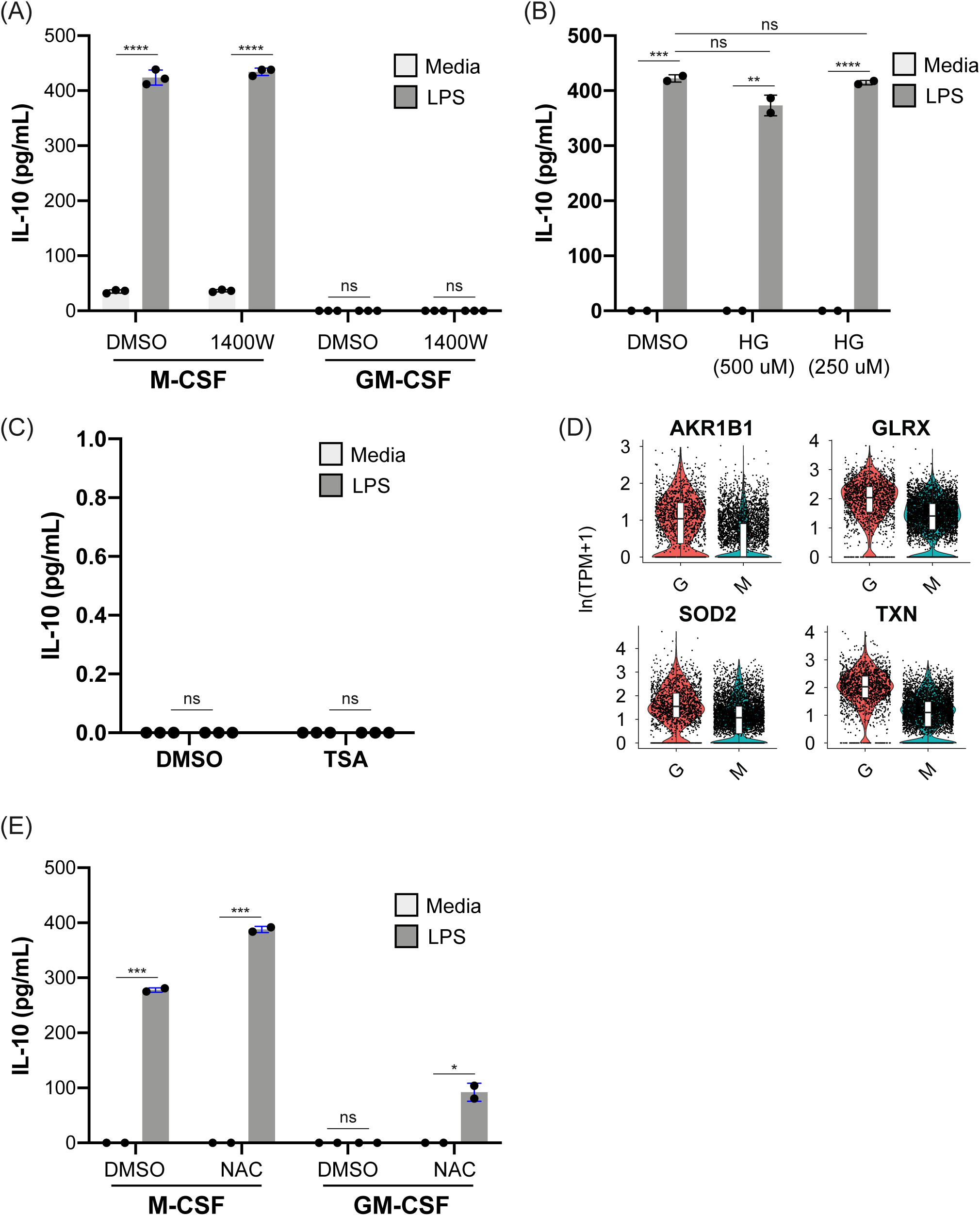
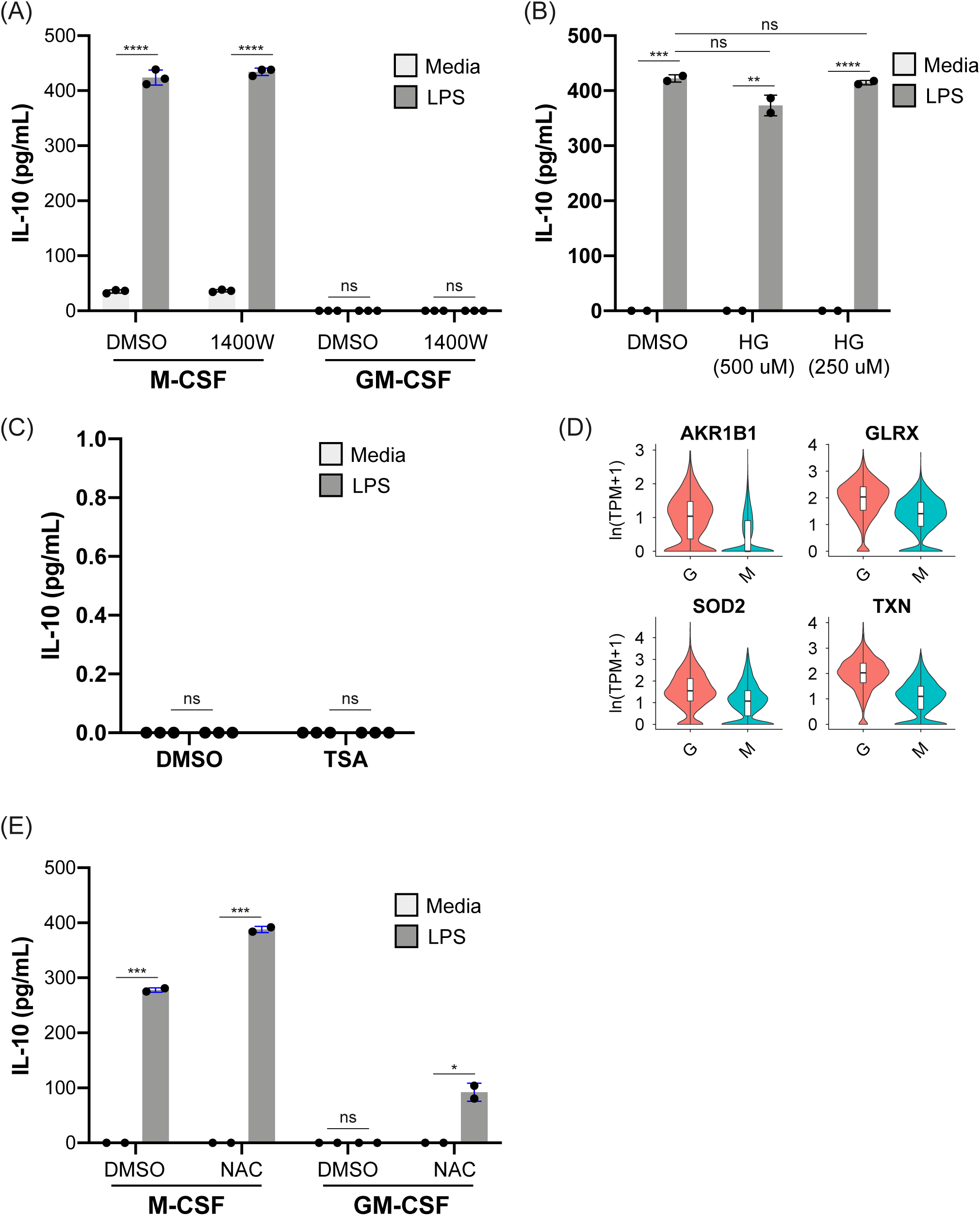
Interference with inducible nitric oxide synthase or salt-inducible kinase activity does not enhance IL-10 production while scavenging oxidative radicals with N-acetylcysteine in MO-GMCSF cells does. (A) Secretion of IL-10 in MO-GMCSF or MO-MCSF cells following incubation with 1400W, an iNOS inhibitor, and treatment with 100 ng/mL LPS stimulation. (****, p < 0.0001, ns = not significant). Representative of 3 donors. (B) Secretion of IL-10 in MO-MCSF cells following incubation with HG-9-91-01, a salt-inducible kinase inhibitor, and treatment with 100 ng/mL LPS stimulation. (****, p < 0.0001, ***, p < 0.001, **, p < 0.01). Representative of 3 donors. (C) Secretion of IL-10 in MO-GMCSF cells that were pre-treated with 100 nM trichostatin-A (TSA) or vehicle control prior to 6-day differentiation and next stimulated with 100 ng/mL LPS for 16 hours. (****, p < 0.0001, ns = not significant). Representative of 2 donors. (D) Log-normalized expression of genes associated with oxidative signaling in MO-MCSF or MO-GMCSF cells. (E) Secretion of IL-10 in MO-GMCSF or MO-MCSF cells that were pre-treated with 5 mM N-acetylcysteine prior to 6-day differentiation and next stimulated 100 ng/mL LPS for 16 hours. (***, p < 0.001, *, p <0.05, ns = not significant). Representative of 3 donors.

Salt inducible kinase (SIK) inhibitors were previously identified as potentiators of IL-10 production in macrophages following their differentiation into macrophages (Lombardi et al., 2016; MacKenzie et al., 2013). We examined the capacity of salt inducible kinase inhibitor, HG-9-91-01, to enhance IL-10 production in MO-MCSF and MO-GMCSF cells. MO-MCSF and MO-GMCSF cells were incubated with HG-9-91-01 for 2 hours prior to the addition of LPS; however, we observed no increase in the production of IL-10 in SIK-inhibitor treated cells relative to the vehicle control (**Figure 4B**). To test the hypothesis that the absence of robust IL-10 production in MO-GMCSF cells was mediated by the activity of epigenetic regulators, we next examined the contribution of Class I and Class II histone deacetylases by pre-treating the cells with trichostatin A (TSA), an inhibitor of these enzymes, prior to the addition of GM-CSF. Pre-treatment with 100 nM TSA had no impact on the capacity of MO-GMCSF cells to produce IL-10 (**Figure 4C**). In parallel, we sought to interrogate the effects of M-CSF and GM-CSF treatment on chromatin accessibility through the lens of ATAC-sequencing with a specific focus on the IL-10 locus. IL-10 did not achieve a statistically significant p-value after Bonferroni hypothesis correction (**Supplemental Table 3**).

Interfering with previously described networks involved in IL-10 production or phenotypic stability in macrophages had minimal impact in enhancing IL-10 production in MO-GMCSF cells.

We next sought to review our unbiased transcriptomic data for additional insight and hypotheses for programs that we could perturb in MO-GMCSF cells to restore IL-10 production. In MO-GMCSF cells, we observed statistically significant enrichment of pathways associated with oxidoreductase signaling including TXN, SOD2, and GLRX (**Figure 4D**). We next sought to examine the contribution of oxidative signaling in MO-GMCSF cell function by first pre-treating monocytes with N-acetylcysteine (NAC) for one hour prior to the addition of M-CSF or GM-CSF. After 6 days, the cells were washed and stimulated with LPS. As expected, production of IL-10 was observed in all MO-MCSF cells. Consistent with a model whereby oxidative signaling contributes to the suppression of IL-10 release in MO-GMCSF cells, pre-treatment of monocytes with NAC prior to the addition of differentiation cytokines mediated an increase in IL-10 release following LPS treatment, though not to levels of MO-MCSF cells (**Figure 4E**). Together, these data demonstrate the utility of unbiased approaches to augment hypothesis generation as well as underscore the contribution of immunometabolic circuits to macrophage state.

## Discussion

Macrophage states, mostly studied through the lens of M1 and M2 activation, have been studied extensively both *in vivo* and *in vitro*. Many models have been proposed to describe the mechanisms that underlie macrophage state in a diverse array of settings (Lawrence and Natoli, 2011; Murray, 2017). Traditionally, macrophages have long been considered to be plastic cells that can polarize to distinct states as needed. However, challenges exist when determining macrophage state and examining phenotypic transitions in tissues in a cell type such as primary human macrophages. Unlike lymphocytes, few, if any, robust binary markers exist for cell state determination in primary human macrophages. Here, we studied a widely utilized model of human macrophages using high-resolution tools to define novel markers of cell state and examine macrophage functional stability. Using our systematic approach to examine macrophage state, we show that MO-GMCSF cells adopt a stable phenotype with respect to IL-10 production and that this phenotype can be disrupted by the addition of scavengers of reactive oxygen species.

We show that conventional surface markers are of limited utility to discern macrophage state using a binary gating strategy. These results are consistent with countless previous studies that utilize analog differences in surface protein expression instead of digital expression as done with lymphocytes (Becher et al., 2014; Roussel et al., 2017; Schulz et al., 2019). Cytokine profiling in response to TLR ligands helped discriminate LPS-tolerized macrophages in both MO-MCSF and MO-GMCSF states. Moreover, cytokine profiling aided in discriminating MO-MCSF and MO-GMCSF cells through the lens of IL-10 production, consistent with previous studies of MO-MCSF and MO-GMCSF cells (Lacey et al., 2012; Sander et al., 2017). Here, our work extends upon these previous studies by examining the temporal dimension of macrophage state and reveals the striking stability in macrophage state that can be achieved.

To address challenges in characterizing macrophage state using protein expression, we conducted scRNAseq of MO-MCSF and MO-GMCSF cells and identified numerous novel markers of macrophage state. Namely, in MO-GMCSF cells, we identified strong enrichment of pathways associated with oxidative signaling consistent with emerging models relating metabolism in driving macrophage state (Cameron et al., 2019; West et al., 2011). We speculate that this interplay between oxidative signaling and metabolic state influence a multitude of macrophage functions, such as bacterial clearance and atherosclerosis, *in vivo* (Di Gioia et al., 2020; Huang et al., 2018; Wang et al., 2020). We identified and validated several novel markers of macrophage state. We also highlight shared programs that are active across multiple models of inflammatory macrophages such as thioredoxin signaling, as well as programs such as ALOX5AP that are specific to differentiation cue. Previous studies in human macrophages are consistent with our unbiased study of macrophage gene expression. Studies by Werz and colleagues similarly observed digital expression of ALOX5AP in MO-GMCSF cells and demonstrate that this digital protein expression is correlated with digital production of the inflammatory lipid PGE2 (Werz et al., 2018). Taken together, these data identify novel markers of macrophage state as well as regulatory axes that should be considered when selecting *in vitro* culture models. PGE2 has been previously demonstrated to modulate macrophage state through the lens of metabolic function and cytokine release highlighting a critical role for this lipid in immunoregulation (MacKenzie et al., 2013; Sanin et al., 2018). It is appealing to hypothesize that high levels of PGE2 production and low IL-10 production may contribute to amplification of an inflammatory macrophage state.

Using IL-10 release as a tool to interrogate functional stability of MO-MCSF and MO-GMCSF cells. Surprisingly, we observed that MO-MCSF cells retained functional plasticity with respect to IL-10 production while MO-GMCSF cells did not. Previous studies in primary human macrophages demonstrate that classical M(LPS+IFN-*γ*) macrophages do not upregulate CD206 and CD200R markers upon treatment with IL-4 whereas M-CSF macrophages do upregulate CD206 and CD200R when treated with IL-4 (Van den Bossche et al., 2016). IL-10 production has traditionally been associated with anti-inflammatory macrophage states. Our *in vitro* results demonstrate a shared property among inflammatory macrophages where limited phenotypic transitions from inflammatory to anti-inflammatory states are possible. Previous studies in M(LPS+IFN-*γ*) human macrophages did not identify a strategy to reverse the inflammatory state. Here, using an unbiased systems-level analysis, we identified oxidative signaling as a driver of IL-10 suppression. Our results demonstrating an association between GM-CSF signaling and reactive oxygen species is consistent with several other studies demonstrating a role for GM-CSF in modulating ROS production in diverse myeloid cell types (Kasahara et al., 2016; Spath et al., 2017; Subramanian Vignesh et al., 2013). Our protein expression analysis revealed that thioredoxin was highly expressed in both MO-GMCSF and M(LPS+IFN-*γ*) cells suggesting a potentially conserved mechanism whereby oxidative signaling contributes to inflammatory macrophage phenotypic stability. Previous work has highlighted the transcriptional regulator BHLHE40 as an essential repressor of IL-10 production in murine cells treated with GM-CSF; however, regulation of BHLHE40 expression in M(LPS+IFN-*γ*) has not been explored (Huynh et al., 2018). Upregulation of BHLHE40 has been observed in response to DNA damaging agents, and recent studies demonstrate that DNA damage mediated by reactive oxygen species is observed in murine macrophages following the addition of inflammatory agents such as LPS and IFN-*γ* (Cameron et al., 2019; Kim et al., 2014). Examining the regulation of BHLHE40 expression and the contribution of BHLHE40 in inflammatory macrophage stability should be the subject of future inquiry.

While many *in vitro* macrophage models focus on driving a particular state in a digital fashion providing a single cytokine in isolation, here, we began to investigate the role of combinations of cytokines in mediating cell state. Again, even when M-CSF is provided in excess, we observed that GM-CSF is incredibly potent in mediating IL-10 suppression. In tissues where GM-CSF is constitutively present like the lungs, it is highly plausible that GM-CSF may play a dominant role in modulating macrophage function independent of ontogeny. In fact, previous studies of isolated murine alveolar macrophages show undetectable or very low levels of IL-10 production in response to TLR ligands (Sajti et al., 2020; Salez et al., 2000). These data raise the possibility that MO-GMCSF cells may provide a functionally similar macrophage to those isolated from tissues consistent with transcriptional analyses performed by Sander; however, additional functional studies are necessary to more fully characterize axes of *in vitro* and tissue-resident macrophage state. These data suggest that complex immune environments may contain dominant soluble mediators of cell state, similar to how GM-CSF drives a low-IL-10 producing state even at low concentrations. How these relationships scale for other cytokine pairs and other responses has been partially investigated in the past; here we build on the experimental designs employed in those studies with a cellular reprogramming model involving staggered addition of cytokines or TLR ligands and suggest future work to explore combinatorial signal perception and gene expression manifolds more exhaustively in macrophages (Natarajan et al., 2006; Xue et al., 2014).

Since human monocyte-derived macrophages have limited to non-existent replicative capacity, leveraging the markers identified in our study may allow recovery of cellular history in a manner akin to fate-mapping experiments conducted in genetically engineered mice. Future studies should further define the relationship between signal inputs and the expression of markers identified in our study. We found that GM-CSF was a driver of ALOX5AP expression; however, it is not known if there are other cytokines that induce ALOX5AP expression. Similarly, we identified TXN as a marker of MO-GMCSF cells as well as MO(LPS+IFN-*γ*) cells. Shared expression of TXN in MO(LPS+IFN-*γ*) and MO-GMCSF cells suggests a shared response network, possibly oxidative signaling, and may serve as an equally valuable marker to inform metabolic history of immune cells. Lastly, as we continue to decipher the relationship between marker expression and cytokine stimulation, it will be essential to understand the relationship between signal duration and magnitude and marker expression to fully exploit the potential of these novel markers.

In conclusion, our study demonstrates the phenotypic stability of MO-GMCSF cells by exploring the temporal dimension of macrophage state. Here, we demonstrate using a combination of immune profiling assays with scRNAseq that oxidative signaling contributes to limited IL-10 release in human macrophages. For future work, we propose deeper study of metabolic and oxidative signaling in both MO-GMCSF and other models of inflammatory macrophages to identify shared mechanisms of phenotypic stability.

## Methods

### PBMC Isolation and Monocyte Differentiation

Deidentified buffy coats from healthy human donors were obtained from Massachusetts General Hospital. PBMCs were isolated from buffy coats by density-based centrifugation using Ficoll (GE Healthcare). CD14+ monocytes were isolated from PBMCs using a CD14 positive-selection kit (Stemcell). Isolated monocytes were differentiated to MO-MCSF or MO-GMCSF in RPMI 1640 (ThermoFisher Scientific) supplemented with 10% heat-inactivated FBS (ThermoFisher Scientific), 1% HEPES, and 1% L-glutamine. Media was further supplemented with either 50 ng/mL M-CSF or 50 ng/mL GM-CSF unless otherwise specified (Biolegend, MCSF: 572902 and GM-CSF: 574802). Monocytes were cultured on low-adhesion tissue culture plates (Corning) for 6 days. Where specified, additional polarization stimuli were added on day 4 (10 ng/mL IFN-*γ*, 10 ng/mL IL-4, 10 ng/mL IL-6, 10 ng/mL IL-10, and 10 ng/mL LPS) for a total of 6 days in culture. For 6 well dishes, cells were seeded at a density of 0.5e6 cells/mL. For 12- and 96-well dishes, cells were seeded at a density of 0.25e6 cells/mL.

### Surface Epitope Staining and Flow Cytometry

Macrophages were detached from low-adhesion tissue culture plates using pre-warmed 1X PBS (Ca2+ and Mg2+ free) (Corning, 21-040-CV) supplemented with 2mM EDTA. Detached macrophages were counted, pelleted at 500xg, and blocked using Human TruStain FcX (Biolegend: 422301) at 2x for 5 min at room temperature. Cells were then stained directly using a pre-mixed antibody panel cocktail for 15 min in the dark at 4° C. Cells were then washed twice with FACS buffer (1X Ca2+ and Mg2+ free PBS, 2% heat-inactivated FBS, and 1mM EDTA) prior to fixation in 2% paraformaldehyde for 15 minutes at room temperature. Following fixation, cells were washed twice with FACS buffer and then analyzed by flow cytometry.

#### Antibody panels are as follows

Panel 1: CD11b-BV421 (Biolegend, 301323), CD32-PECy7 (Biolegend, 303213), CD36-APC (Biolegend, 336207), HLA-DR-BV510 (Biolegend, 307645)

Panel 2: CD11b-BV421 (Biolegend, 301323), CD274-PE (Biolegend, 329705), CD86-APC (Biolegend, 305411), CD64-BV510 (Biolegend, 305027)

Additional antibodies utilized in this study included CD36-PE (Biolegend, 336205), CD206-PE (Biolegend, 321105), CD123 (Biolegend, 306005), CD125 (R&D Systems, FAB253P), CD115-BV421 (Biolegend, 347321), CD116-PE (Biolegend, 305908).

Samples were analyzed on a BD LSR Fortessa.

### Intracellular Cytokine Staining and Flow Cytometry

MO-MCSF cells were simultaneously treated with Brefeldin A (Biolegend, 420601) and stimulated with 100 ng/mL LPS for 12 hours. After this incubation, cells were detached from low-adhesion tissue culture plates using pre-warmed 1X PBS (Ca2+ and Mg2+ free) (Corning, 21-040-CV) supplemented with 2mM EDTA. Detached macrophages were counted, pelleted at 500xg, and resuspended in Fixation Buffer (Biolegend, 420801) and allowed to fix for 20 minutes at room temperature in the dark. Following fixation, cells were pelleted at 500xg and resuspended in Intracellular Staining Perm Wash Buffer (Biolegend, 421002). Three total washes in Intracellular Staining Perm Wash Buffer were performed. Following permeabilization, cells were pelleted for 5 minutes at 500xg and resuspended in 50 µL Intracellular Staining Perm Wash Buffer supplemented with 5 µL of TruStain FcX (Biolegend, 422302) for 10 minutes at room temperature. Then, 50 µL Intracellular Staining Perm Wash Buffer supplemented with 5 µL of IL-10-BV421 antibody (Biolegend, 501421) was added for an additional 20 minutes at room temperature. Following antibody incubation, cells were pelleted for 5 minutes at 500xg and washed twice with Intracellular Staining Perm Wash Buffer prior to resuspension in FACS buffer (1X Ca2+ and Mg2+ free PBS, 2% heat inactivated FBS, and 1mM EDTA) prior to analysis on a BD LSR Fortessa.

### Luminex

MO-GMCSF or MO-MCSF cells were treated for 16 hours with either 100 ng/mL LPS or 100 ng/mL Pam3CSK4 prior to analysis of cytokine release. Multiplexed cytokine measurements on macrophage cell culture supernatants were performed using a custom Bio-Plex Express Human Cytokine kit (Bio-Rad 17004073) according to manufacturer’s instructions, using a Bio-Plex 3D Suspension Array System (Bio-Rad). Measured cytokines included IL-6, IFN-*γ*, GM-CSF, TNF*α*, IL-1*β*, IP-10 (CXCL10), MCP-1 (CCL-2), IL-8, IL-10, and IL-12p70. Supernatants were stored at -80°C until analysis.

### Enzyme-linked Immunosorbent Assays (ELISA)

0.25e6 cells were plated per well of a 12 well dish for cytokine release assays. Cells were stimulated with either 100 ng/mL LPS or 100 ng/mL Pam3Csk4 for 16 hours. Supernatants were analyzed fresh or stored at -80°C until analysis. Cytokine release was measured by ELISA using TNF*α*, IL-6, and IL-10 ELISA kits (Biolegend: TNF*α* 430201, IL-6 430501, and IL-10 430601).

### Single Cell RNA Sequencing

We performed SeqWell analysis as previously described (Gierahn et al., 2017). Briefly, beads were loaded into the PDMS array followed by subsequent addition of ∼12,000 cells. After periodic gentle rocking and washing, a polycarbonate membrane was attached to the array surface.

After membrane attachment, the array was transferred to a dish containing cell lysis buffer. After lysis, buffer was exchanged for hybridization buffer to enable RNA hybridization to barcoded beads. Beads were removed from microwell arrays by scraping the array with a microscope slide.

Recovered beads were transferred to a 1.5 mL tube and reverse transcription was performed using Maxima H-reverse transcriptase (Invitrogen). Beads were then treated with ExoI to digest unoccupied primer sites (NEB).

cDNA was amplified and the pooled PCR library was purified, and tagmentation was subsequently performed. Libraries were sequenced on an Illumina NextSeq500. 20 bases were allocated (12 bp cell barcode and 8 bp UMI) for Read 1, which was primed using Custom Read 1 Primer.

### Transcriptome Alignment

Read alignment was performed as in described previously (Macosko et al., 2015). Briefly, for each NextSeq sequencing run, raw sequencing data was converted to FASTQ files using bcl2fastq2 that were demultiplexed by Nextera N700 indices corresponding to individual samples. Reads were first aligned to HgRC19 and individual reads were tagged according to the 12-bp barcode sequence and the 8-bp UMI contained in read 1 of each fragment. Following alignment reads were binned and collapsed onto 12-bp cell barcodes that correspond to individual beads using DropSeq tools (http://mccarrolllab.com/dropseq/). Barcodes were collapsed with a single-base error tolerance (Hamming distance =1) with additional provision for single insertions or deletions. An identical collapsing scheme (Hamming distance = 1) was then applied to unique molecular identifiers to obtain quantitative counts of individual mRNA molecules.

### Single Cell RNAseq Data Pre-Processing

We identified a total of 1,742 good-quality GM-CSF-macrophages (donor 1: 399, donor 2: 1,343) and 4,803 good-quality M-CSF-treated cells (donor 1: 1,941, donor 2: 2,862) with total transcripts >2,500 and <75,000 and mitochondrial read percentage less than 20%. After filtering based on quality control metrics, raw count matrices were normalized by library size. For each cell, UMI-collapsed gene expression counts were divided by total transcript counts for that cell, multiplied by a scale factor of 10,000, then natural logarithm-normalized using Seurat v3.0.2. Log-transformed expression levels were standard scaled using Seurat’s ScaleData function. During scaling, technical covariates (percentage mitochondrial reads and number of UMIs) were regressed out using a generalized linear model, as implemented in Seurat, and the residuals of the generalized linear model were used as ‘corrected’ gene expression values for downstream scaling.

### Differential Expression Analysis

Differential gene expression between MO-MCSF and MO-GMCSF macrophages was assessed by Wilcoxon and AUC tests using Seurat’s FindMarkers function and log-transformed expression levels [ln(TPM+1)]. Genes with absolute log2 fold change > 0.25 and Bonferroni-corrected p-values < 0.05 were considered differentially expressed genes (DEGs). DEGs with AUC < 0.7 or AUC > 0.3 were selected as markers of MO-GMCSF and MO-MCSF macrophages, respectively. Differential expression analyses were conducted on individual donors separately, as well as on all cells together.

### Gene Ontology Enrichment Analysis

Gene symbols were mapped to Entrez gene ids using org.Hs.eg.db annotations. GO enrichment was analyzed by hypergeometric test using the clusterProfiler package’s enrichGO function against a background set of genes captured in all sequencing runs with > 100 total counts across runs. GO terms with Benjamini-Hochberg-correct p-values < 0.05 were considered significantly enriched. Significant GO terms were collapsed into representative GO terms using clusterProfiler’s simplify function and a similarity threshold of 0.7.

## Data availability

All RNA-seq data are available in GEO under accession number XXXX. Code needed to complete the RNA-seq analysis can be found on github.com/cgunnars/mcsf-gmcsf-sc

### Western Blotting

Macrophages were lysed using RIPA buffer with Proteinase Inhibitor. Protein concentration was estimated using a BCA assay. 15ug of protein were loaded per sample in a Bolt 4-12% Bis-Tris Plus Gel. PageRuler Plus was used as a molecular weight ladder. The gel was transferred to a nitrocellulose membrane using the iBlot 2 dry blotting system (Invitrogen). Membranes were blocked for 1 hour using Odyssey blocking buffer at room temperature. Membranes were next incubated in primary antibody shaking overnight at 4C. TRX (Santa Cruz, sc-166393), *β*-actin (Biolegend, Poly6221), ALOX5AP (GeneTex, GTX106689), and *β*-actin (Santa Cruz, sc-47778) were diluted with TRIS buffered saline containing Tween-20 (TBST). After incubation with the primary antibody, membranes were washed three times with TBST and then incubated with a fluorescent secondary antibody (LiCor IRDye 680 or LiCor IRDye 800) at a dilution of 1:10,000 in TBST for 1 hour. The membranes were then washed three times with TBST. The western blots were imaged by Odyssey CLx.

### STAT3 Phosphorylation Analysis

MO-MCSF or MO-GMCSF cells were washed once with PBS following 6 days of differentiation, and then stimulated with media containing vehicle or media containing 10 or 50 ng/mL IL-10 for 15 minutes. Following incubation with IL-10, cells were washed once with cold PBS and lysed immediately on ice. Western blotting was performed as described above. Membranes were incubated with either a total STAT3 antibody (Cell Signaling Technology, Clone: 79D7) or a phosphorylated STAT3 antibody (Cell Signaling Technology, Clone: D3A7).

### Macrophage Durability Assay

To evaluate the stability of TLR-mediated IL-10 release by macrophages, following 6 days of differentiation with M-CSF or GM-CSF, MO-MCSF cells or MO-GMCSF cells were washed with pre-warmed media without cytokines and then incubated with media supplemented with 50 ng/mL M-CSF, 50 ng/mL GM-CSF or no cytokine. Following 3 days of incubation, cells were washed with PBS and then stimulated with 100 ng/mL LPS for 16 hours prior to quantification of IL-10 release by ELISA.

### GM-CSF + M-CSF Checkerboard Assay

For the GM-CSF/M-CSF checkerboard assay, a range of concentrations of GM-CSF and M-CSF (0, 0.5 ng/mL, 5 ng/mL, and 50 ng/mL) were premixed in a pairwise fashion prior to the addition of monocytes. Monocytes were then added to the individual wells to commence differentiation. Monocytes were allowed to differentiate for 6 days. After 6 days, cells were washed twice with PBS and incubated with maintenance media with or without 100 ng/mL LPS.

### GM-CSF and Drug Perturbation Experiments

To evaluate the impact of inhibition of iNOS on IL-10 production in MO-MCSF and MO-GMCSF cells, monocytes were pretreated with 1400W, an iNOS inhibitor, at a final concentration of 100 μM for 1 hour prior to the addition of 50 ng/mL GM-CSF or 50 ng/mL M-CSF. 1400W was dissolved in DMSO. After 6 days of differentiation, cells were washed and then treated with 100 ng/mL LPS for 16 hours. Cytokine release was compared to cells treated with DMSO alone (0.1% v/v final concentration). Supernatants were analyzed fresh or stored at -80°C until analysis.

To evaluate the capacity of salt-inducible kinase inhibitor or PGE2 treatment on production of IL-10 by MO-MCSF cells, cells were pre-treated with DMSO (0.1% v/v), 250 or 500 nM HG-9-91-01 (VWR, dissolved in DMSO, final concentration 0.1% v/v DMSO), or 5 or 10 μM PGE2 (Santa Cruz Biotechnology) for two hours prior to treatment with LPS. Without washing, cells were then treated with 100 ng/mL LPS for 16 hours prior to analysis by ELISA.

To evaluate the impact of inhibiting class I and II histone deacetylases, monocytes were pretreated with 100 nM trichostatin A (TSA, Sigma Aldrich). Monocytes were pre-treated for 1 hour prior to the addition of 50 ng/mL GM-CSF or 50 ng/mL M-CSF to induce differentiation. After 6 days of differentiation, cells were washed and then treated with 100 ng/mL LPS for 16 hours. Supernatants were analyzed fresh or stored at -80°C until analysis.

To evaluate the impact of scavenging oxidative radicals prior to differentiation, monocytes were pretreated with 5 mM N-acetylcysteine (NAC) which was dissolved directly in cell culture media and filtered through 0.2 μm filter. Monocytes were pre-treated for 1 hour prior to the addition of 50 ng/mL GM-CSF or 50 ng/mL M-CSF to induce differentiation. After 6 days of differentiation, cells were washed and then treated with 100 ng/mL LPS for 16 hours. Supernatants were analyzed fresh or stored at -80°C until analysis.

### ATAC Sequencing and Analysis

MO-GMCSF and MO-MCSF cells were prepared for ATAC sequencing as previously described (Buenrostro et al., 2015). Briefly, cells were detached and washed twice in PBS and diluted to a final concentration of 1e6 cells/mL. 5,000 cells were then mixed with TD buffer, 10% NP-40, 20x Illumina Tn5, and water. The reaction was allowed to proceed for 30 mins with shaking at 37°C with mixing. Transposed DNA is isolated using a PCR purification kit (Qiagen) and eluted. 5 cycles of PCR were performed on the transposed product with indexing primers and qPCR was performed on 5 µL of the PCR product to identify the optimal number of total cycles for each sample. The remaining PCR mix was stored on ice until the optimal number of additional cycles was identified. The amplified PCR product was purified using a PCR purification kit (Qiagen), and DNA concentration was determined using a PhiX standard curve. DNA from each sample was pooled at approximately equal input and sequenced on an Illumina NextSeq 500.

Raw sequencing data was converted to FASTQ files using bcl2fastq2 that were demultiplexed according to ATACseq sample indices corresponding to individual samples. Reads were aligned to GRCh38.

Chromatin accessibility near transcriptional start sites was analyzed following the standards set forth by the Encode Consortium (Buenrostro et al., 2015). Briefly, scores were obtained by aligning ATAC reads to genome and performing peak calling and next examining peak summit heights at +/- 1kb around the transcriptional start site (TSS) of each gene, then averaging peak scores with a linearly decreasing weight with increasing distance to TSS.

## Acknowledgments

This work was supported by MIT Biological Engineering (BDB), The Ragon Institute of MGH, MIT, and Harvard (AKS, BDB, SMF), NIH CFAR P30 AI060354, The Bill and Melinda Gates Foundation (AKS, SMF), NIH R01 AI123286 (SMF), NIH T32AI132120-01 (GHB), NIH R01 AI12328 (SMF, GHB), NIH R01 GM081871 (BLH, BB)Searle Scholars Program (AKS), Beckman Young Investigator Program (AKS), Pew-Stewart Scholars Program for Cancer Research (AKS), and the Sloan Fellowship in Chemistry (AKS). CYI was funded by an HHMI Medical Research Fellowship. CG is supported by an NSF Graduate Research Fellowship. B.H. is supported by the Department of Defense (DoD) through the National Defense Science and Engineering Graduate Fellowship (NDSEG).

## Author Information Contributions

CYI, CG, GHB, AN, NAA, MHW, TKH, SLS, BH, BB, AKS, SMF, and BDB conceived and performed experiments. CYI, CG, GHB, AN, NAA, MHW, TKH, SLS, BH, BB, AKS, SMF, and BDB all performed data analysis. All authors contributed to writing of the manuscript and analysis of experimental results.

## Supplemental Figure Legends

**Supplemental Figure 1: Analysis of macrophage surface marker expression using a digital gating strategy to identify the percentage of macrophages expressing a given marker (MO-GMCSF cells)**. Representative of 2 donors.

**Supplemental Figure 2: Analysis of macrophage surface marker expression using an analog quantification strategy (mean fluorescence intensity, MFI) to quantify the abundance of a given marker on the macrophage surface (MO-GMCSF cells)**. Representative of 2 donors.

**Supplemental Figure 3: Quantification of CD274 across multiple donors reveals upregulation of CD274 following IFN-γ or LPS stimulation (MO-GMCSF cells)**. Each colored line is an individual donor.

**Supplemental Figure 4: Analysis of IL-10 production in MO-MCSF cells following LPS stimulation using intracellular cytokine staining and flow cytometry**. Production of IL-10 following 100 ng/mL LPS stimulation was monitored using intracellular cytokine staining.

**Supplemental Figure 5: Read mapping quality in samples**. Violin plots depict number of transcripts (A), genes (B), and percentage of reads aligning to mitochondrial genes (C), separated by GM-CSF and M-CSF treatment and donor.

**Supplemental Table 1: Differential expression of all genes captured in scRNAseq analysis of MO-GMCSF and MO-MCSF cells**. The AUC column presents the results of the area under the curve analysis used to identify marker genes for MO-GMCSF and MO-MCSF cells. The avg_log2FC column presents the log2(fold change) in gene expression for MO-GMCSF cells relative to MO-MCSF cells. The power column represents the classification power of a particular gene. The pct.g column indicates the percentage of MO-GMCSF cells expressing a particular gene. The pct.m column indicates the percentage of MO-MCSF cells expressing a particular gene. The p_val column represents the p-value of differential gene expression between MO-GMCSF and MO-MCSF cells as determined by Wilcoxon rank sums test. The p_val_adj column represents the p-value of differential gene expression between MO-GMCSF and MO-MCSF cells following Benjamini-Hochberg correction.

**Supplemental Table 2: Gene ontology analysis of marker genes from analysis of MO-GMCSF and MO-MCSF cells by scRNAseq**. The ID column represents the ID of the gene ontology term. The Description column describes and summarizes the gene ontology term. The GeneRatio column captures the number of significant marker genes belonging to a particular gene ontology category out of all the significant marker genes for MO-GMCSF or MO-MCSF cells. The BgRatio column captures the number of captured genes belonging to a particular gene ontology category out of all captured genes for MO-GMCSF or MO-MCSF cells. The background set of genes was constructed by identifying genes captured in all sequencing runs with >100 total counts across runs. The pvalue column captures the results of the hypergeometric test. The p.adjust column is after filtering gene ontology terms by semantic similarity (Benjamini-Hochberg-corrected p-value < 0.05; semantic similarity threshold = 0.5). The qvalue column was determined using a false-discovery rate approach. The geneID column captures the gene names of those genes belonging to the particular Gene Ontology term. The Count column enumerates the number of significant marker genes falling into a particular Gene Ontology category.

**Supplemental Table 3: Chromatin accessibility scores of MO-MCSF and MO-GMCSF cells**. The Gene column represents the gene ID. The GM-CSF Score column summarizes the chromatin accessibility scores for all 6 MO-GMCSF ATACseq samples. The M-CSF Score column summarizes the chromatin accessibility scores for all 6 MO-MCSF ATACseq samples. The Mean GM-CSF Score column presents the mean chromatin accessibility score for all 6 MO-GMCSF samples. The Mean GM-CSF Score column presents the mean chromatin accessibility score for all 6 MO-GMCSF samples. The Mean M-CSF Score column presents the mean chromatin accessibility score for all 6 MO-MCSF samples. The Uncorrected p-value column is the result of a t-test for each gene.

## References

Becher, B., Schlitzer, A., Chen, J., Mair, F., Sumatoh, H.R., Teng, K.W.W., Low, D., Ruedl, C., Riccardi-Castagnoli, P., Poidinger, M., et al. (2014). High-dimensional analysis of the murine myeloid cell system. Nat. Immunol. 15, 1181–1189.

Biswas, S.K., and Mantovani, A. (2010). Macrophage plasticity and interaction with lymphocyte subsets: cancer as a paradigm. Nat. Immunol. 11, 889–896.

Bogdan, C. (2001). Nitric oxide and the immune response. Nat. Immunol. 2, 907–916.

Buenrostro, J., Wu, B., Chang, H., and Greenleaf, W. (2015). ATAC-seq: A Method for Assaying Chromatin Accessibility Genome-Wide. Curr. Protoc. Mol. Biol. Ed. Frederick M Ausubel Al 109, 21.29.1-21.29.9.

Cameron, A.M., Castoldi, A., Sanin, D.E., Flachsmann, L.J., Field, C.S., Puleston, D.J., Kyle, R.L., Patterson, A.E., Hässler, F., Buescher, J.M., et al. (2019). Inflammatory macrophage dependence on NAD + salvage is a consequence of reactive oxygen species–mediated DNA damage. Nat. Immunol. 20, 420.

Di Gioia, M., Spreafico, R., Springstead, J.R., Mendelson, M.M., Joehanes, R., Levy, D., and Zanoni, I. (2020). Endogenous oxidized phospholipids reprogram cellular metabolism and boost hyperinflammation. Nat. Immunol. 21, 42–53.

DuPage, M., and Bluestone, J.A. (2016). Harnessing the plasticity of CD4 + T cells to treat immune-mediated disease. Nat. Rev. Immunol. 16, 149–163.

Fleetwood, A.J., Lawrence, T., Hamilton, J.A., and Cook, A.D. (2007). Granulocyte-macrophage colony-stimulating factor (CSF) and macrophage CSF-dependent macrophage phenotypes display differences in cytokine profiles and transcription factor activities: implications for CSF blockade in inflammation. J. Immunol. Baltim. Md 1950 178, 5245–5252.

Gierahn, T.M., Wadsworth Ii, M.H., Hughes, T.K., Bryson, B.D., Butler, A., Satija, R., Fortune, S., Love, J.C., and Shalek, A.K. (2017). Seq-Well: portable, low-cost RNA sequencing of single cells at high throughput. Nat. Methods 14, 395–398.

Hamilton, J.A. (2019). GM-CSF-Dependent Inflammatory Pathways. Front. Immunol.10.

Huang, L., Nazarova, E.V., Tan, S., Liu, Y., and Russell, D.G. (2018). Growth of Mycobacterium tuberculosis in vivo segregates with host macrophage metabolism and ontogeny. J. Exp. Med. 215, 1135–1152.

Huynh, J.P., Lin, C.-C., Kimmey, J.M., Jarjour, N.N., Schwarzkopf, E.A., Bradstreet, T.R., Shchukina, I., Shpynov, O., Weaver, C.T., Taneja, R., et al. (2018). Bhlhe40 is an essential repressor of IL-10 during Mycobacterium tuberculosis infection. J. Exp. Med. 215, 1823–1838.

Kasahara, S., Jhingran, A., Dhingra, S., Salem, A., Cramer, R.A., and Hohl, T.M. (2016). Role of Granulocyte-Macrophage Colony-Stimulating Factor Signaling in Regulating Neutrophil Antifungal Activity and the Oxidative Burst During Respiratory Fungal Challenge. J. Infect. Dis. 213, 1289–1298.

Kim, J., D’Annibale, S., Magliozzi, R., Low, T.Y., Jansen, P., Shaltiel, I.A., Mohammed, S., Heck, A.J.R., Medema, R.H., and Guardavaccaro, D. (2014). USP17-and SCFβTrCP-Regulated Degradation of DEC1 Controls the DNA Damage Response. Mol. Cell. Biol. 34, 4177–4185.

Lacey, D.C., Achuthan, A., Fleetwood, A.J., Dinh, H., Roiniotis, J., Scholz, G.M., Chang, M.W., Beckman, S.K., Cook, A.D., and Hamilton, J.A. (2012). Defining GM-CSF– and Macrophage-CSF–Dependent Macrophage Responses by In Vitro Models. J. Immunol. 188, 5752–5765.

Lavin, Y., Winter, D., Blecher-Gonen, R., David, E., Keren-Shaul, H., Merad, M., Jung, S., and Amit, I. (2014). Tissue-Resident Macrophage Enhancer Landscapes Are Shaped by the Local Microenvironment. Cell 159, 1312–1326.

Lawrence, T., and Natoli, G. (2011). Transcriptional regulation of macrophage polarization: enabling diversity with identity. Nat. Rev. Immunol. 11, 750–761.

Lee, S., Kim, S.M., and Lee, R.T. (2013). Thioredoxin and thioredoxin target proteins: from molecular mechanisms to functional significance. Antioxid. Redox Signal. 18, 1165–1207.

Liu, S.X., Gustafson, H.H., Jackson, D.L., Pun, S.H., and Trapnell, C. (2020). Trajectory analysis quantifies transcriptional plasticity during macrophage polarization. Sci. Rep. 10.

Liu, Z., Gu, Y., Chakarov, S., Bleriot, C., Kwok, I., Chen, X., Shin, A., Huang, W., Dress, R.J., Dutertre, C.-A., et al. (2019). Fate Mapping via Ms4a3-Expression History Traces Monocyte-Derived Cells. Cell 178, 1509-1525.e19.

Locati, M., Curtale, G., and Mantovani, A. (2020). Diversity, Mechanisms, and Significance of Macrophage Plasticity. Annu. Rev. Pathol. Mech. Dis. 15, 123–147.

Lombardi, M.S., Gilliéron, C., Dietrich, D., and Gabay, C. (2016). SIK inhibition in human myeloid cells modulates TLR and IL-1R signaling and induces an antiinflammatory phenotype. J. Leukoc. Biol. 99, 711–721.

MacKenzie, K.F., Clark, K., Naqvi, S., McGuire, V.A., Nöehren, G., Kristariyanto, Y., Bosch, M. van den, Mudaliar, M., McCarthy, P.C., Pattison, M.J., et al. (2013). PGE2 Induces Macrophage IL-10 Production and a Regulatory-like Phenotype via a Protein Kinase A–SIK–CRTC3 Pathway. J. Immunol. 190, 565–577.

Macosko, E.Z., Basu, A., Satija, R., Nemesh, J., Shekhar, K., Goldman, M., Tirosh, I., Bialas, A.R., Kamitaki, N., Martersteck, E.M., et al. (2015). Highly Parallel Genome-wide Expression Profiling of Individual Cells Using Nanoliter Droplets. Cell 161, 1202–1214.

Murray, P.J. (2017). Macrophage Polarization. Annu. Rev. Physiol. 79, 541–566.

Murray, P.J., Allen, J.E., Biswas, S.K., Fisher, E.A., Gilroy, D.W., Goerdt, S., Gordon, S., Hamilton, J.A., Ivashkiv, L.B., Lawrence, T., et al. (2014). Macrophage Activation and Polarization: Nomenclature and Experimental Guidelines. Immunity 41, 14–20.

Natarajan, M., Lin, K.-M., Hsueh, R.C., Sternweis, P.C., and Ranganathan, R. (2006). A global analysis of cross-talk in a mammalian cellular signalling network. Nat. Cell Biol. 8, 571–580.

Niemand, C., Nimmesgern, A., Haan, S., Fischer, P., Schaper, F., Rossaint, R., Heinrich, P.C., and Müller-Newen, G. (2003). Activation of STAT3 by IL-6 and IL-10 in Primary Human Macrophages Is Differentially Modulated by Suppressor of Cytokine Signaling 3. J. Immunol. 170, 3263–3272.

Plugis, N.M., Weng, N., Zhao, Q., Palanski, B.A., Maecker, H.T., Habtezion, A., and Khosla, C. (2018). Interleukin 4 is inactivated via selective disulfide-bond reduction by extracellular thioredoxin. Proc. Natl. Acad. Sci. 201805288.

Roussel, M., Ferrell, P.B., Greenplate, A.R., Lhomme, F., Gallou, S.L., Diggins, K.E., Johnson, D.B., and Irish, J.M. (2017). Mass cytometry deep phenotyping of human mononuclear phagocytes and myeloid-derived suppressor cells from human blood and bone marrow. J. Leukoc. Biol. 102, 437–447.

Sajti, E., Link, V.M., Ouyang, Z., Spann, N.J., Westin, E., Romanoski, C.E., Fonseca, G.J., Prince, L.S., and Glass, C.K. (2020). Transcriptomic and epigenetic mechanisms underlying myeloid diversity in the lung. Nat. Immunol. 21, 221–231.

Salez, L., Singer, M., Balloy, V., Créminon, C., and Chignard, M. (2000). Lack of IL-10 synthesis by murine alveolar macrophages upon lipopolysaccharide exposure. Comparison with peritoneal macrophages. J. Leukoc. Biol. 67, 545–552.

Sander, J., Schmidt, S.V., Cirovic, B., McGovern, N., Papantonopoulou, O., Hardt, A.-L., Aschenbrenner, A.C., Kreer, C., Quast, T., Xu, A.M., et al. (2017). Cellular Differentiation of Human Monocytes Is Regulated by Time-Dependent Interleukin-4 Signaling and the Transcriptional Regulator NCOR2. Immunity 47, 1051-1066.e12.

Sanin, D.E., Matsushita, M., Klein Geltink, R.I., Grzes, K.M., van Teijlingen Bakker, N., Corrado, M., Kabat, A.M., Buck, M.D., Qiu, J., Lawless, S.J., et al. (2018). Mitochondrial Membrane Potential Regulates Nuclear Gene Expression in Macrophages Exposed to Prostaglandin E2. Immunity 49, 1021-1033.e6.

Schneemann, M., and Schoedon, G. (2002). Species differences in macrophage NO production are important. Nat. Immunol. 3, 102–102.

Schulz, D., Severin, Y., Zanotelli, V.R.T., and Bodenmiller, B. (2019). In-Depth Characterization of Monocyte-Derived Macrophages using a Mass Cytometry-Based Phagocytosis Assay. Sci. Rep. 9, 1–12.

Spath, S., Komuczki, J., Hermann, M., Pelczar, P., Mair, F., Schreiner, B., and Becher, B. (2017). Dysregulation of the Cytokine GM-CSF Induces Spontaneous Phagocyte Invasion and Immunopathology in the Central Nervous System. Immunity 46, 245–260.

Subramanian Vignesh, K., Landero Figueroa, J.A., Porollo, A., Caruso, J.A., and Deepe, G.S. (2013). Granulocyte Macrophage-Colony Stimulating Factor Induced Zn Sequestration Enhances Macrophage Superoxide and Limits Intracellular Pathogen Survival. Immunity 39, 697–710.

Van den Bossche, J., Baardman, J., Otto, N.A., van der Velden, S., Neele, A.E., van den Berg, S.M., Luque-Martin, R., Chen, H.-J., Boshuizen, M.C.S., Ahmed, M., et al. (2016). Mitochondrial Dysfunction Prevents Repolarization of Inflammatory Macrophages. Cell Rep. 17, 684–696.

Wang, Y., Ji, N., Gong, X., Ni, S., Xu, L., and Zhang, H. (2020). Thioredoxin-1 attenuates atherosclerosis development through inhibiting NLRP3 inflammasome. Endocrine.

Werz, O., Gerstmeier, J., Libreros, S., Rosa, X.D. la, Werner, M., Norris, P.C., Chiang, N., and Serhan, C.N. (2018). Human macrophages differentially produce specific resolvin or leukotriene signals that depend on bacterial pathogenicity. Nat. Commun. 9, 1–12.

West, A.P., Brodsky, I.E., Rahner, C., Woo, D.K., Erdjument-Bromage, H., Tempst, P., Walsh, M.C., Choi, Y., Shadel, G.S., and Ghosh, S. (2011). TLR signalling augments macrophage bactericidal activity through mitochondrial ROS. Nature 472, 476–480.

Xue, J., Schmidt, S.V., Sander, J., Draffehn, A., Krebs, W., Quester, I., De Nardo, D., Gohel, T.D., Emde, M., Schmidleithner, L., et al. (2014). Transcriptome-Based Network Analysis Reveals a Spectrum Model of Human Macrophage Activation. Immunity 40, 274–288.

