## Supplemental Figures for "GM-CSF differentiation of human monocytes stabilizes macrophage state via oxidative signaling" (from another version of this preprint: version 1, 2020-10-01)

#### Supplemental Figure 1

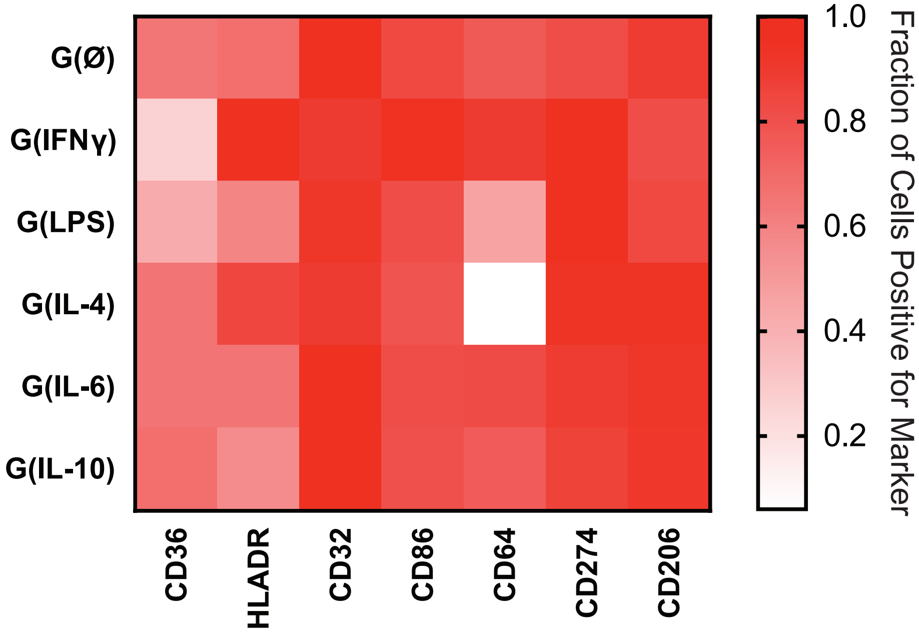

**Supplemental Figure 2**

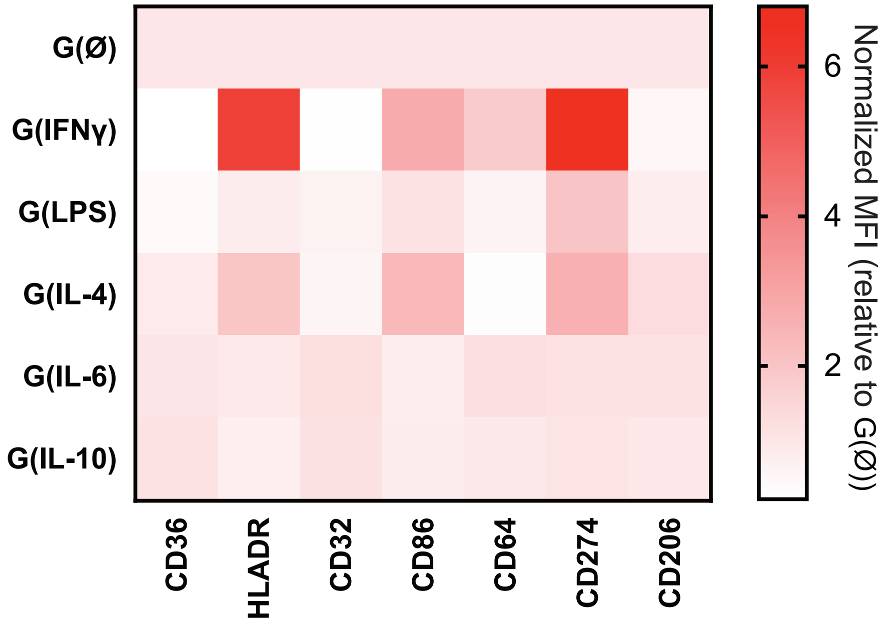

**Supplemental Figure 3**

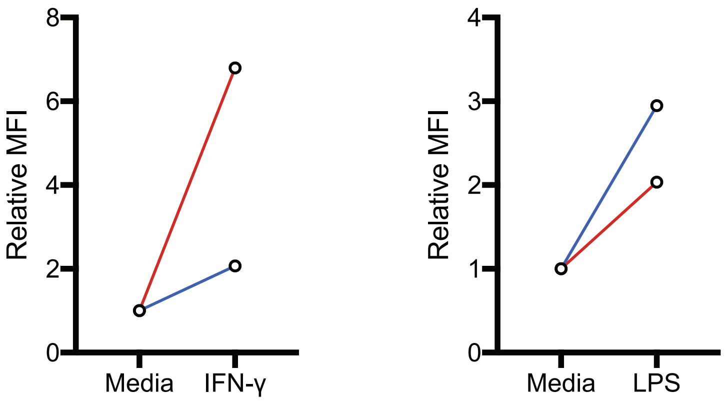

### Supplemental Figure 4

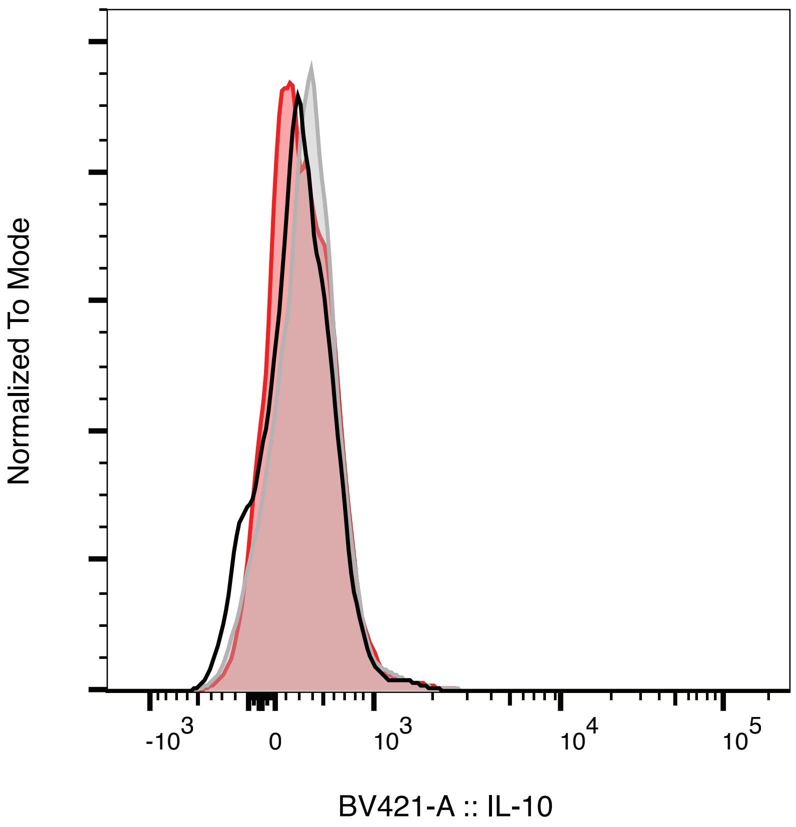


| | Sample Name | Subset Name | Count |
|---|---|---|---|
| ☐ | Test_Unstim_Unstained.fcs | Single Cells | 7974 |
| ☐ | Test_Unstim_IL10_BV421.fcs | Single Cells | 7939 |
| ☐ | Test_LPS_IL10_BV421.fcs | Single Cells | 6376 |

### Supplemental Figure 5

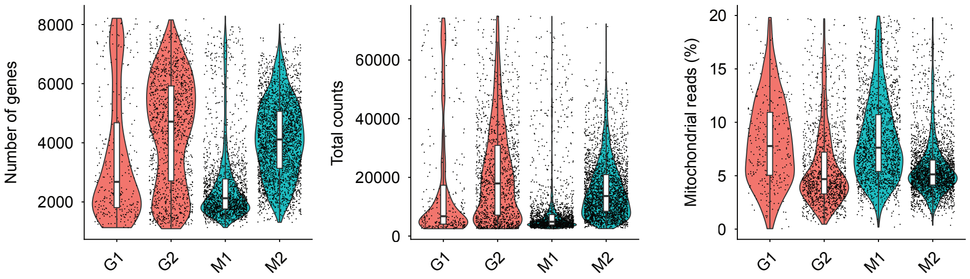
